# Modification of the microstructure of the CERN- CLEAR-VHEE beam at the picosecond scale modifies ZFE morphogenesis but has no impact on hydrogen peroxide production

**DOI:** 10.1101/2024.12.19.629203

**Authors:** Houda Kacem, Louis Kunz, Pierre Korysko, Jonathan Ollivier, Pelagia Tsoutsou, Adrien Martinotti, Vilde Rieker, Joseph Bateman, Wilfrid Farabolini, Gérard Baldacchino, Billy W. Loo, Charles L. Limoli, Manjit Dosanjh, Roberto Corsini, Marie-Catherine Vozenin

## Abstract

FLASH has emerged as a significant breakthrough for the future of radiation oncology, as it reduces complications while preserving the tumor killing efficacy. To define the beam parameters for future clinical translation, Very High Energy Electrons (VHEE) delivered at CLEAR and able to reach deep seated tumors were used in conjunction with a FLASH-validated Intermediate Energy Electron (IIE) beam and a 160-225 keV X-ray beam, collectively able to deliver dose rates spanning from 1 Gy/min to 10^11^ Gy/s. High-throughput chemical assays were used to investigate radiochemical effects of FLASH, while zebrafish embryos served as a model to evaluate its impact on biological outcomes and morphogenesis. This study is the first comprehensive exploration investigating the impact of a large range of dose rates and various temporal parameters from early physico-chemical events to a complex biological system. Data derived at CLEAR revealed that the intensity of the bunch is a critical factor for observing the sparing effect of FLASH and uncovered an unforeseen biological response when electrons are delivered over the picosecond timescale. Present data also suggests that scanning with high intensity beamlets will be optimal for the future clinical translation of FLASH.

**Highlights:** To investigate the physics parameters required to trigger the FLASH sparing effect, CLEAR/VHEE/CERN beam macro/microstructure was varied. We show that delivery at the picosecond scale:

- reduces alteration in the morphogenesis of zebrafish embryos, but
- has no impact on secondary hydrogen peroxide production,

**Graphical abstract:** 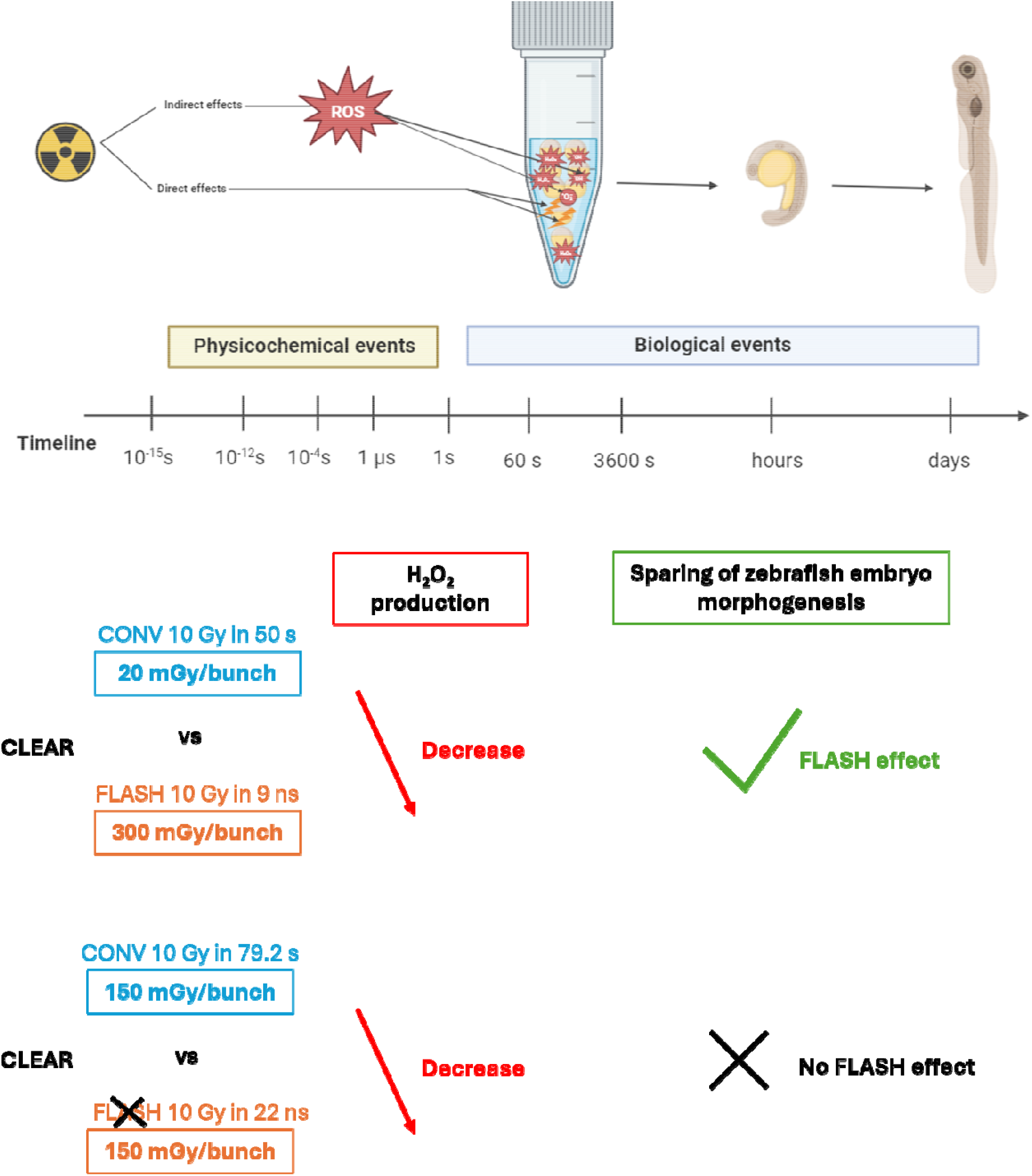

## 1. Introduction

Radiotherapy at ultra-high dose rate (UHDR), also known as FLASH radiotherapy, is currently seen as one of the most promising innovations in radiation oncology [1]. Experimental evidence from various preclinical models indicated that shortening radiation exposure times below 200 milliseconds can protect normal tissues while remaining efficient against tumors, a phenomenon referred to as the FLASH effect. Recent studies have investigated the biological basis involved in normal tissue sparing [2,3]. Data reveal that the pathological response typically activated in normal tissue by radiotherapy at standard dose rates (≤ 1 Gy/min) is not triggered by FLASH [2,3]. Low levels of inflammation and protection of normal cells, including vascular, epithelial, mesenchymal and stem cells are consistently found after FLASH exposure whereas tumor cells are killed equally [3]. More recent studies suggest that FLASH remains effective under conditions classically known to induce tumor radiation resistance such as hypoxia [4] and immuno-depletion [5]. These findings are promising and could expand the therapeutic window of radiotherapy, offering new opportunities for cancer cure. However, several critical questions remain for safe and meaningful clinical transfer. While understanding the mechanisms responsible for the differential effect between normal versus tumor tissues would be significant, the priority to unlock the full potential of FLASH radiotherapy for the improvement of cancer treatment and patient well-being is to define the beam characteristics required to trigger the FLASH effect [3].

To date, the FLASH effect has been observed to be particle and beam-type agnostic. It has been reported with various beams, including intermediate energy pulsed electron (4-10 MeV) [6–8], photon [9,10] superconducting devices [11,12], and modified clinical proton beams [13–15]. Nevertheless, the precise parameters able to trigger the FLASH effect remain uncertain. While the average dose rate of 40 Gy/s is often cited as the reference FLASH dose rate, negative reports have been published using this questionable threshold [16–18].

FLASH has opened a new and intriguing area of research aimed at understanding how the duration of dose delivery impacts the biological response to irradiation. Key questions include understanding the temporal dynamics involved in the interaction of energetic electrons and tissues as well as the basis of the differential response triggered between healthy and tumor tissues. While time scales of elementary processes underpinning molecular physics and chemistry are well-known, our initial hypothesis was that the energy transfer from ionizing radiation to the milieu at femtosecond time scales could modify the early physico-chemical events of the water radiolysis cascade by altering the free radical cascades and recombination reactions that begin at sub-picosecond scales and last over milliseconds [19]. We postulated that these events might subsequently modify the downstream biological response and that measurements in cell free aqueous based systems could be used as a surrogate to probe the physical parameters required to trigger the FLASH effect.

The current study was designed to probe this hypothesis and define the temporal parameters required to trigger the FLASH effect using CLEAR at CERN a unique 190 − 210 MeV facility. Very High Energy Electron beam (VHEE), the FLASH-validated 5.5 MeV Intermediate Energy Electron beam (IEE) eRT6/Oriatron beams were used as well as a conventional dose rate with 160 − 225 keV X-ray beam as a reference. The research delved into the physico-chemical and biological effects of these three distinct beams, each characterized by unique temporal structures and covering a broad dose rate spectrum from 1 Gy/min to 10^11^Gy/s, using water, plasmid DNA and zebrafish embryos. Distinctive features specific to each beam-type and temporal structure were identified and showed that the intensity of the bunch delivered over picoseconds at CLEAR and of the pulse delivered over microseconds at eRT6 are critical parameters for preserving ZFE morphogenesis.

## 2. Materials and Methods

### 2.1. Irradiation beam lines

In this study we used a variety of beam lines able to deliver a wide range of dose rates.

1. CLEAR (CERN): 190 − 210 MeV electrons were delivered at conventional 0.125 Gy/s and 0.2 Gy/s) and FLASH (10^8^ Gy/s to 10^11^ Gy/s) dose rates as previously described [20]. Chemical samples and ZFE were placed in individual PCR tubes (0.2 mL) positioned in a 3D-printed holder within a water tank. The samples were then placed in the beam using c-robot [21].
2. eRT6 (Oriatron, PMB/Alcen): 5.5 MeV electrons were delivered at conventional (0.1 Gy/s) and FLASH (60 Gy/s to 10^7^ Gy/s) dose rates as described before [22,23]. Chemical samples and ZFE were placed in individual Eppendorf tubes (2 mL) and positioned vertically in a water tank. CLEAR beam is made of trains, each made up of Gaussian-shaped bunches of 3 ps that can be adjusted. The dose rate can be varied from 0.1 up to 10^11^ Gy/s. eRT6 is made of pulses, each made up of Gaussian-shaped bunches of 4.63 ps that cannot be modified. The dose rate can be varied from 0.1 up to 10^7^ Gy/s. The historical nomenclature was retained, referring to the beam structures as ‘train’ for CLEAR and ‘pulse’ for eRT6, even though they essentially represent the macrostructure.
3. The Xrad 225CX/225 keV (Pxi Precision X-ray) and RS2000 160 keV (Rad Source Technologies) X-ray tubes were used as a reference for photon irradiation at conventional dose rates (0.037 and 0.07 Gy/s). Chemical samples and ZFE were placed in individual Eppendorf tubes (2 mL) and positioned vertically to the X-ray source. Irradiation parameters are detailed in the Supplementary Tables. Dosimetry was performed according to previously cross-validated protocols [24,25]. Both beams were linear accelerators (linacs) that have radiofrequency (RF) cavities but the thermionic injector for eRT6 did not allow to control the bunch structure so, the smallest accessible temporal resolution was the pulse while the CLEAR beam composed of trains of bunches, allowed for the smallest controllable structure to be the individual bunch.

### 2.2. Water radiolysis

#### 2.2.1. G°(H_2_O_2_): Primary yields of hydrogen peroxide

To determine the primary yield of hydrogen peroxide produced during water radiolysis, we followed the production of H_2_O_2_ as a function of HO scavenger concentration (NaNO_2_/NaNO_3_), as described in [26]. Milli-Q water with a resistivity of 18.2 M.Ω.cm was equilibrated to 1 % O_2_ using a hypoxia hood (Biospherix, Xvivo-system X3) and irradiated as indicated in (Table S1, S2&3a, supplementary data). Water samples were immediately probed post-irradiation with Amplex Red assay kit (Thermo Fisher). Fluorescence quantification was performed using Promega Glo-Max plate reader (Excitation: 520 nm; Emission: 580 − 640 nm).

#### 2.2.2. Production of hydrogen peroxide

To get closer to biological conditions, water samples, subject to various conditions known to influence radiation response in biological systems, were used. Oxygen levels, temperature, and scavengers varied according to (Table S12, supplementary data), and water samples were irradiated as previously described (Table S1, S2 &S3a, supplementary data). H_2_O_2_ analysis was conducted similarly to the earlier method.

### 2.3. Simulation method

Chemsimul software was used to calculate the concentration of hydrogen peroxide produced after the homogenous phase of chemistry and across the different dose rates. The software utilizes a set system of differential equations to solve, which requires the use of known primary yields at the start of the simulation. More details can be found on the software website: https://chemsimul.dk/.

### 2.4. DNA damage in pBR322 plasmid

To investigate DNA damage, pBR322 plasmid (Thermo Fisher Inc.) was purified and diluted at 40 ng/μL in deionized, RNAase and DNAase-free water (UltraPure). Plasmids were then irradiated according to the methods described previously in (Table S1, S2&3b, supplementary data). Radiation induced modifications were resolved using Agarose Gel Electrophoresis (AGE) with 0.8 % agarose in Tris-acetate-EDTA (TAE) buffer. The compact supercoiled form and the two relaxed forms, open circular, and linear, were quantified using densitometric analysis (UVITEC Cambridge) and ImageJ software “gels” add-on. DNA damage was investigated in various environmental conditions such as different oxygen levels, scavengers, and labile iron (Fe^2+^) concentrations, as shown in (Table S13, supplementary data). The McMahon model was applied to the data to describe the damage on plasmid resulting from irradiation [27]. The model helped in quantifying the rates of single strand breaks (SSB) and double strand breaks (DSB), denoted as β_S_ and β_D_, respectively. This allowed for a quantitative comparison of the amount of damaged plasmids by different irradiation types and in different environments.

### 2.5. Zebrafish embryo irradiation

AB Wild Type ZF (Danio rerio, #1175, F7 generation, EZRC) were bred to produce zebrafish embryos at the PTZ (CHUV/UNIL, Lausanne, Switzerland). According to Swiss and European ethics regulations, no ethical approval is required for use of ZFE before 5 days of development that were considered as biodosimeters. 4 hours post fertilization (hpf) ZFE were irradiated in water at 28°C using the various beam lines as described in (Table S1, S2&3c, supplementary data). Radiation-induced alterations in ZFE length were measured at 5 days post-fertilization (dpf) following embryo fixation (4% PFA) and microscopic imaging (Evos XL Core Cell Imaging System, Thermo Fisher). Analysis was conducted using ImageJ software. The % ZFE Length Deficit is the following:

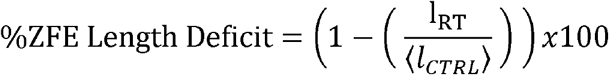

with l_RT_ the length of the irradiated embryo and ⟨l_CTRL_⟩ the average length of the control (non-irradiated) embryos.

### 2.6. Statistical Analysis

Statistical analyses were carried out using GraphPad Prism (v9.1) for water radiolysis, plasmids, and zebrafish embryos (ZFE) experiments. Fluorescence ratio of H_2_O_2_ production was assessed by t-test. For plasmids, the error bars correspond to standard deviation. For ZFE, data are presented as mean +/− SEM and analysis was done using Kruskal-Wallis test.

## 3. Results

### 3.1. G° values of H_2_O_2_ are beam-type and dose-rate independent

Measurement of primary yield of H_2_O_2_ in hypoxia (1% O_2_) was done according to (Table S1, S2&S3a, supplementary data). H_2_O_2_ was used as a surrogate of the initial radiation-induced free radical production. For each concentration of HO^·^ scavenger used, we plotted the G-value against the cubic root of the HO^·^ scavenger following the Swroski’s method [28] (Fig. 1a), and also considered the scavenging capacity [29] (Fig. S1a, b&c, supplementary data).

**Fig. 1.**
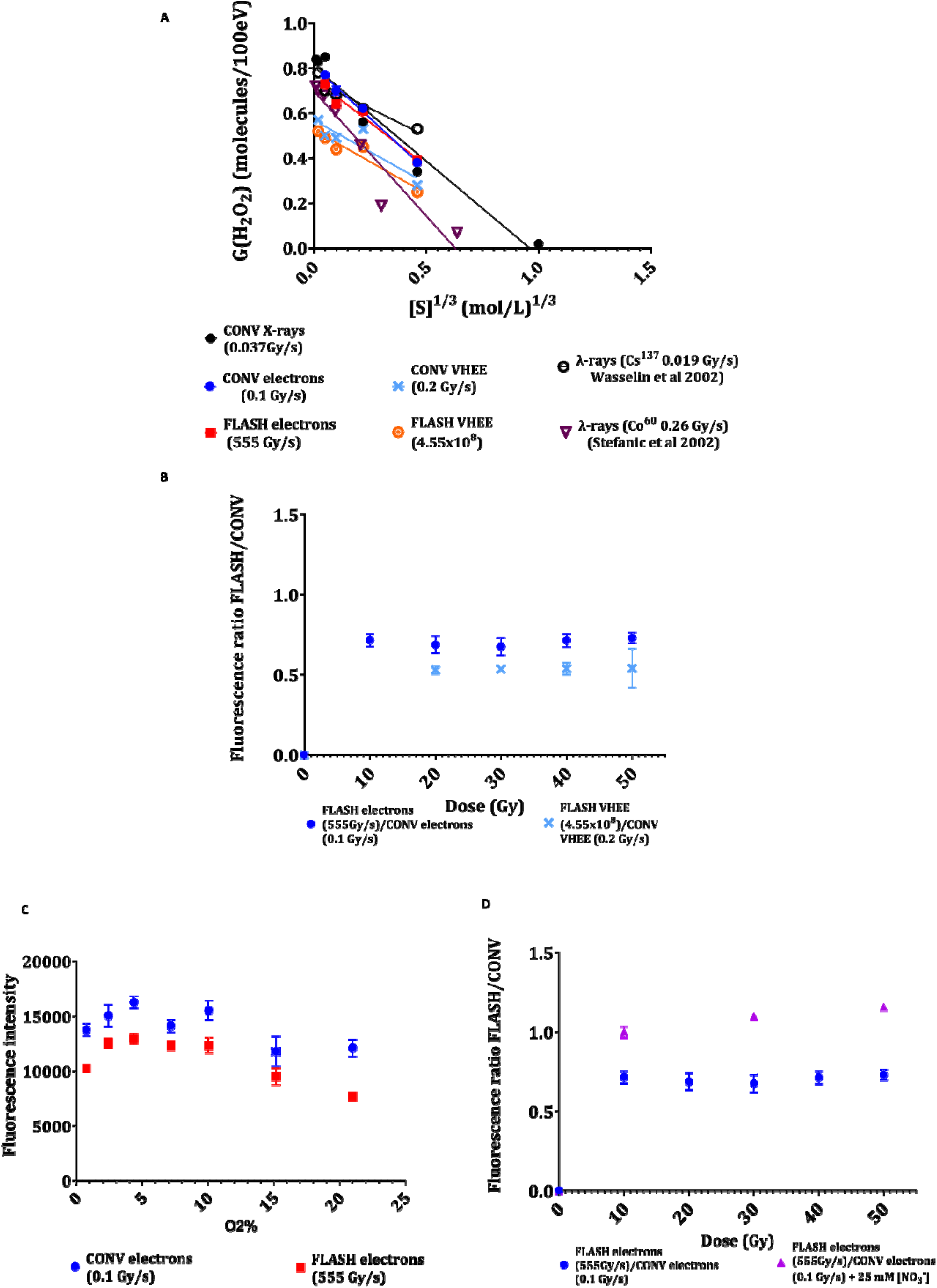
Primary yields of hydrogen peroxide and hydrogen peroxide production after CONV and FLASH irradiation. **A** G-values measured in anoxic conditions (1%O_2_) versus cubic root of a scavenger denoted by [S] (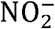 ions for this work) obtained after irradiation with CONV and FLASH (190 - 220 MeV) VHEE, (5.5 MeV) IEE and CONV (225 kVp) X-rays compared to reported experiments performed with CONV (0.6 MeV) ^137^Cs g-rays; [S] = 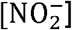 and CONV (1.2 MeV) ^60^Co g-rays; [S] = [Br^-^]. **B** Fluorescence ratio FLASH/CONV of H_2_O_2_ production vs dose at 21%O_2_ in pure water after irradiation with CONV and FLASH VHEE and IEE **C** Fluorescence intensity of H_2_O_2_ production at 30 Gy vs different O_2_ levels after irradiation with CONV and FLASH IEE. **D** Fluorescence ratio FLASH/CONV of H_2_O_2_ production in presence or absence of 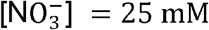 (scavenger of aqueous electrons) after irradiation with CONV and FLASH IEE. Scavenging experiments are results of 6 experiments for IEE and duplicate experiments for CONV X-rays and VHEE (CONV and FLASH). Production of H_2_O_2_ at 21%O_2_ are results of triplicate experiments with both beams. Production of H_2_O_2_ at different O_2_ levels are results of duplicate experiments with IEE. 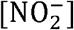 experiments were done with CONV and FLASH IEE. Each experiment had 8 points of measurements per dose.

These values, detailed in (Table 1), were similar for both FLASH and CONV exposures, consistent with the past publications (0.6 − 0.8 molecules/100 eV) [29–31] (Table S4, supplementary data).

**Table 1.** Primary yields of hydrogen peroxide. G°(H_2_O_2_) in hypoxia (1%O_2_) calculated from the Swroski [28] model for the different beam sources.

| Source | Energy | LET<br>[keV/μm] | Dose<br>rate<br>[Gy/s] | [O<sub>2</sub>]<br>[M] | Chemical<br>system | G°(H<sub>2</sub>O<sub>2</sub>)<br>[molecules/100eV] |
| --- | --- | --- | --- | --- | --- | --- |
| CONV<br>VHEE | 190-210<br>MeV | 0.2 | 0.2 | 1.19x10<sup>-5</sup> | [NO<sub>2</sub><sup>-</sup>]<br>/[NO<sub>3</sub><sup>-</sup>] | 0.6 ± 0.02 |
| FLASH<br>VHEE |  |  | 4.55x10<sup>8</sup> |  |  | 0.59 ± 0.02 |
| CONV IEE | 5.5 MeV | 0.2 | 0.1 |  |  | 0.8 ± 0.02 |
| FLASH<br>IEE |  |  | ≥555 |  |  | 0.75 ± 0.03 |
| CONV X-rays | 225 kVp | 3 | 0.037 |  |  | 0.78 ± 0.04 |

### 3.2. FLASH decreases production of hydrogen peroxide

The production of H_2_O_2_ was assessed minutes after water irradiation under atmospheric oxygen conditions (21% O_2_), with or without scavengers. Interestingly, a reduction in H_2_O_2_ production was observed after FLASH compared to CONV exposures, independently of beam type (as shown Fig. 1b). The fluorescence ratio FLASH/CONV dose rate was 0.7 for IEE and 0.5 for VHEE. This suggests that either FLASH diminishes radical production under atmospheric conditions or that H_2_O_2_ is attacked by radicals in overlapping tracks. The complexity of the irradiated solution was then modified to mimic more closely the cellular biochemical conditions. For all dose rates, increasing oxygen concentration increased H_2_O_2_ production in a bell-shaped manner, with a low production found in hypoxic conditions, followed by a peak at physioxic levels (4% O_2_), then decreasing again when approaching atmospheric conditions (Fig. 1c). This effect was independent of the dose rate, but FLASH irradiation reduced H_2_O_2_ production by 17 to 36 % after 30 Gy irradiation (Fig. 1c). The fluorescence reduction after different irradiation doses is shown in (Fig. S2, supplementary data). When nitrate ions [25 mM] were added to scavenge hydrated electrons, H_2_O_2_ production became similar between both modalities and the fluorescence ratio reached 1. (Fig. 1d). Temperature variation, from room temperature (22°C) to physiological human body temperature (37− 38°C), increased H_2_O_2_ production at higher temperatures, and the fluorescence ratio FLASH/CONV was enhanced at 37°C and reached (0.77 − 0.80) (Fig. S3, supplementary data). These results indicate a reduced production of free radicals following FLASH exposure mediated by hydrated electrons, a difference further enhanced at physiological temperature.

### 3.3. Simulated protracted yields of H_2_O_2_ are inconsistent with experimental data

Simulations were then performed using Chemsimul®, a software which utilizes differential equation systems to model the radiolysis of pure water. The equation system implemented is well described in the literature [32]. Protracted concentrations, and particularly those of H_2_O_2_ were also calculated as a function of temperature and by considering the linear energy transfer of MeV-electrons [32]. It is important to note that the primary yields used in simulations were assumed to be constant across the various simulated dose rates. In this way, dose rate effects were only simulated from the homogeneous chemistry stage [19]. Radical reactions and their corresponding rate constants are compiled in (Table S5, supplementary data). Parameters such as pulses, trains, dose rates, pulse duration and total dose were adjusted to closely match the irradiation condition of each water radiolysis experiment. The computed ratios of H_2_O_2_ FLASH/CONV as a function of the dose are provided in (Table 2).

**Table 2.** Simulated concentration ratio of hydrogen peroxide. Simulated [H_2_O_2_] in atmospheric oxygen condition (21%O_2_) with Chemsimul software using IEE and VHEE CONV and FLASH irradiation conditions. The calculation step size is dynamically adjusted according to the rate of concentration. Changes small steps are used during rapid system evolution, typically at the start, while larger steps are employed as the system stabilizes and evolves more slowly. This adaptive step sizing ensures that the simulations are computationally efficient, minimizing both memory usage and processing time. Chemsimul requires the system to be homogeneous at the onset of the simulation, so we utilized known primary yields to model the protracted production of H_2_O_2_, beyond the homogenous phase of chemistry, and across various dose rates. Consequently, dose rate effects were simulated beginning from the homogeneous chemistry stage [19].

| Model | $[O_2]$<br>[M] | Dose [Gy] | $\frac{[H_2O_2]_{FLASH\ IEE}}{[H_2O_2]_{CONV\ IEE}}$ | $\frac{[H_2O_2]_{FLASH\ VHEE}}{[H_2O_2]_{CONV\ VHEE}}$ |
| --- | --- | --- | --- | --- |
| Chemsimul<sup>®</sup> | $2.50 \times 10^{-4}$ | 10 | 1.51 | 1.20 |
|  |  | 20 | 1.55 | 1.17 |
|  |  | 30 | 1.63 | 1.20 |
|  |  | 40 | 1.73 | 1.23 |
|  |  | 50 | 1.84 | 1.27 |

Ratios ranged from 1.51 − 1.84 and 1.17 − 1.27 for IEE and VHEE respectively. The calculated H_2_O_2_ concentration was higher for FLASH compared to CONV, contradicting experimental data. Simulations in presence of 25 mM nitrate ions and/or at physiological temperature conditions (37_°_, C) enhanced ratio up to 1.57 − 1.72 and 1.65 − 1.79 for IEE respectively as shown in (Table S6 a&b, supplementary data). This discrepancy highlights an unexpected behavior arising under FLASH conditions [33] or the limitations of the simulations used.

### 3.4. Plasmid DNA damage is dose rate insensitive

To move a step closer to biological systems, plasmid irradiation was undertaken using CONV X-rays, IEE and VHEE CONV and FLASH according to (Table S1, S2 &S3b, supplementary data). DNA damage was investigated under various conditions. In atmospheric conditions, with the plasmid in pure water, a dose-dependent induction of single strand breaks (SSB) and double strand breaks (DSB) was found but this was dose-rate independent (Fig. 2a). The summary of the results of DSB (β_D_) and SSB yields (β_S_) calculated at atmospheric oxygen conditions produced by these systematic investigations are compiled in (Table 3).

**Table 3.**
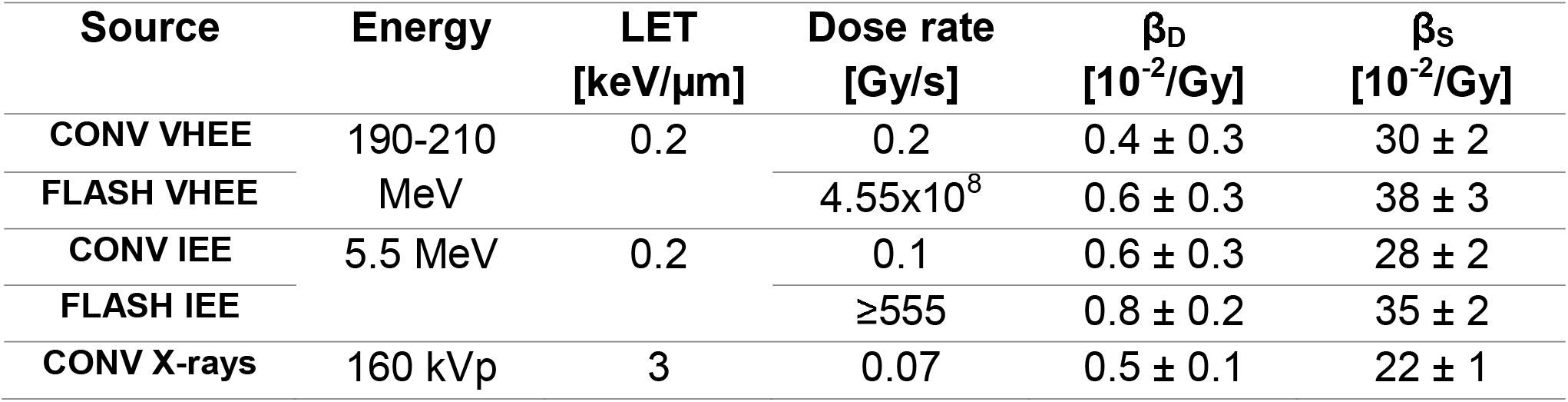
Summary table of DSB (β_D_) and SSB yields (β_s_). DNA damage yield quantification at atmospheric oxygen conditions from different beam structures.

**Fig. 2.**
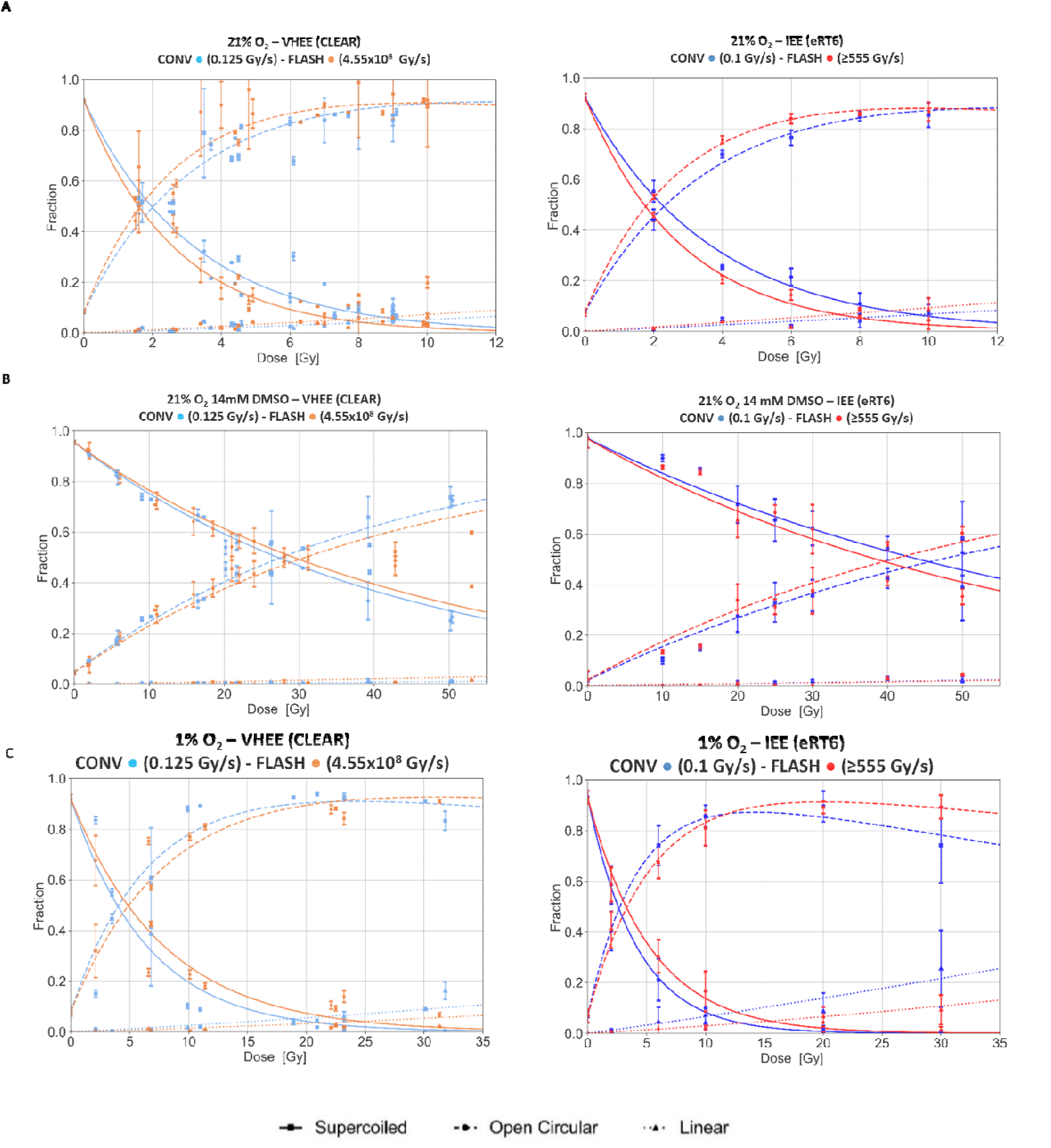
DNA damage in plasmids. **A** at **B** In presence of DMSO and **C** At with CONV and FLASH VHEE and IEE beams. Results are from three independent experiments for all the beams.

Adding 14 mM Dimethylsulfoxide (DMSO), a concentration 10 times lower than cellular antioxidant levels, significantly protected the plasmid from radiolytic damage, with a dose modifying factor (DMF) of about 10-fold which was also dose rate independent (Fig. 2b). In addition, under hypoxic conditions a damage reduction with DMF of about 2-fold is observed, due to the radiosensitizer effect of oxygen. Under hypoxia and increased Fe2+concentrations, used to mimic the tumor environment, DNA damage remained dose-rate independent (Fig. 2c and Fig. S4, supplementary data).

### 3.5. Intense bunches at CLEAR drive the FLASH sparing effect in Zebrafish embryos

Ultimately, we irradiated ZFE with the various beams and parameters (Fig. 3) and compared these biological results with the chemistry results described above.

**Fig. 3.**
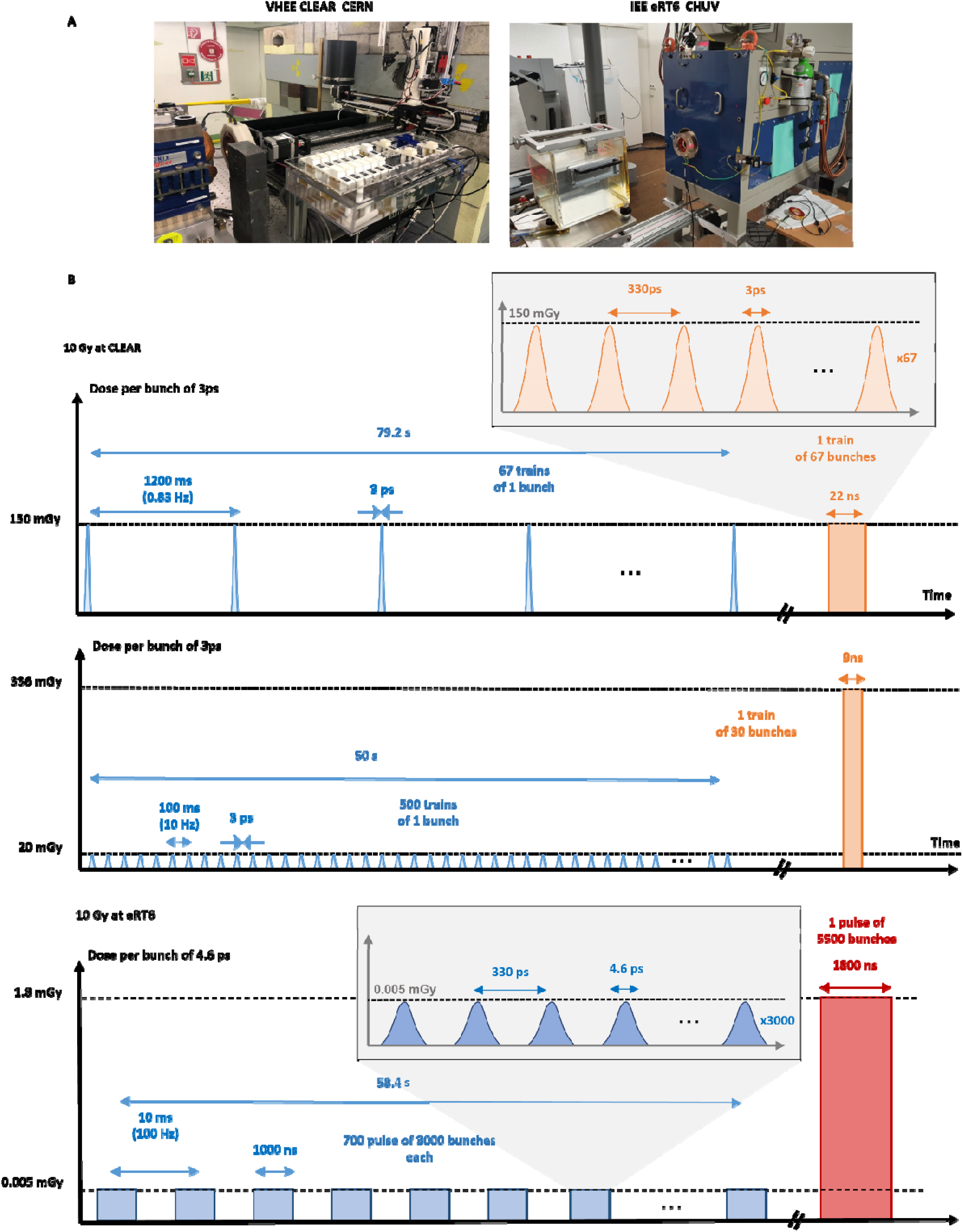
Description of CLEAR (VHEE) and eRT6 (IEE) beam structure. **A** Experimental set up; **B** An example of beam structure after 10 Gy irradiation, orange and red representation are used for FLASH modality and blue is used for conventional modality. VHEE: very high energy electrons and IEE: intermediate energy electrons.

ZFE irradiation was performed using X-rays, IEE and VHEE at conventional and ultra-high dose rates. X-rays at 0.037 Gy/s and IEE at 0.1 Gy/s produced similar dose-dependent defects in the morphogenesis of ZFE, with a length deficit at 10 Gy (D10) reaching 50 % with X-rays and 45 % with IEE compared to non-irradiated controls. FLASH IEE delivered in one single pulse of 10 Gy (Table S2 and S3c, supplementary data), the D10 was only 30 %, showing the FLASH sparing effect and a DMF of ∼1.5 (Fig. 4a).

**Fig. 4.**
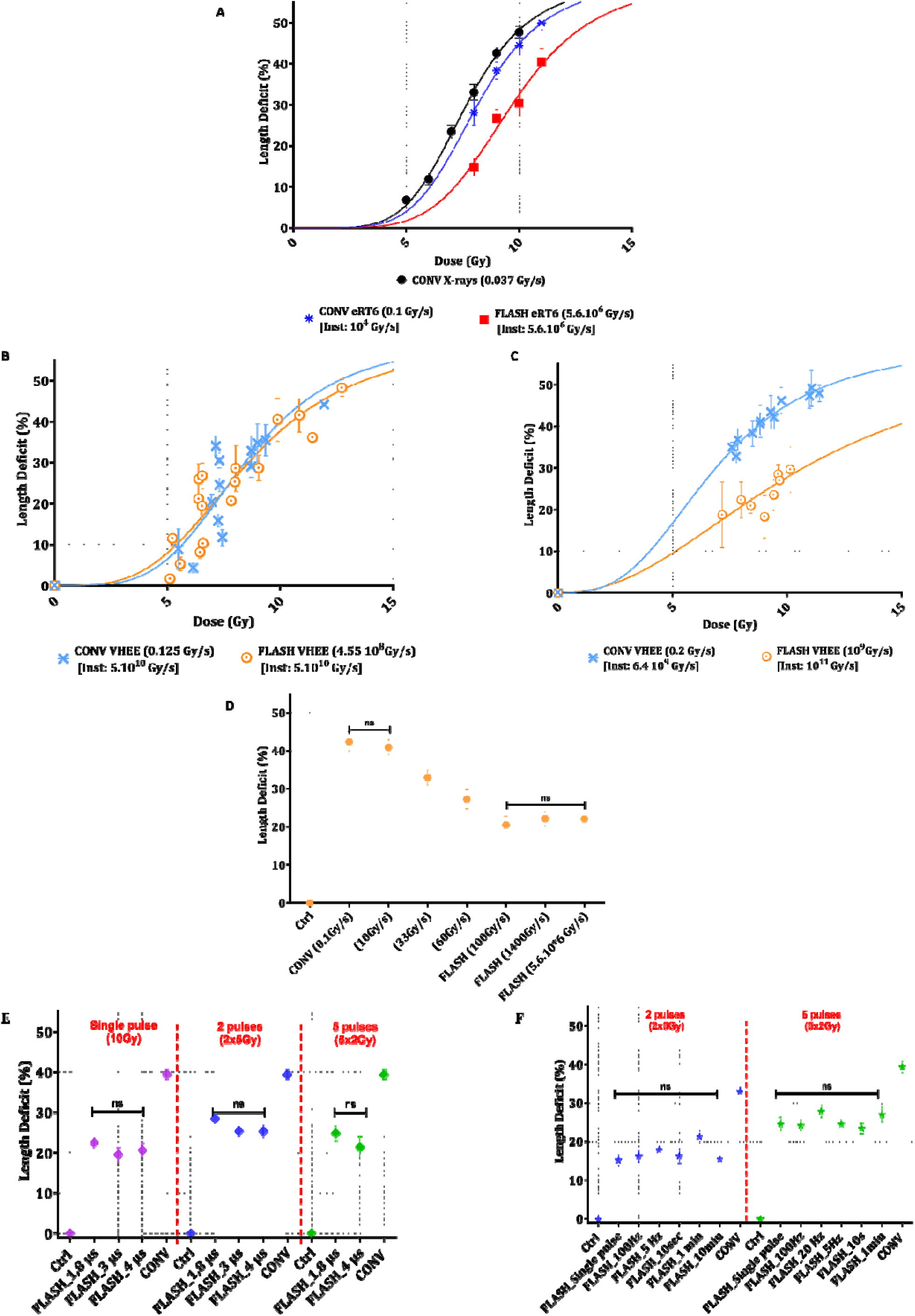
Parametrization experiments on 4hpf ZFE. **A** Evaluation of length deficiency and dose response of 4hpf ZFE after exposure to X-rays (0.037 Gy/s), CONV (0.1 Gy/s) and FLASH (5.6x106 Gy/s) IEE. **B** Evaluation of length deficiency and dose response of 4hpf ZFE after exposure to CONV 0.125 Gy/s and FLASH 4.88x108 Gy/s VHEE with a similar dose rate in the bunch =5x1010 Gy/s. **C** Evaluation of length deficiency and dose response of 4hpf ZFE after exposure to VHEE delivered CONV 0.2 Gy/s and dose rate in the bunch=6.4x10 Gy/s and FLASH 10 Gy/s) and dose rate in the bunch= 1011 Gy/s respectively. **D** Evaluation of length deficiency and dose response of 4hpf ZFE after exposure to increasing dose rates **E** Different pulse widths and **F** Different frequencies. For IEE, embryos (n = 20) per dose and for VHEE irradiations, embryos (n = 8) per dose. Results are the average of 3 experiments for IEE (CONV and FLASH) and 2 experiments for CONV X-rays and VHEE (CONV and FLASH).

With CLEAR-VHEE, when the total dose was delivered with an average dose rate of 0.125 Gy/s versus 4.55x10^8^ Gy/s but using a similar dose in the bunch of 150 mGy, and a similar dose rate in the bunch of 5x10^10^Gy/s, no differential effect was observed. The length deficit reached 40% in both cases falling between the length deficit curves obtained with X-rays/IEE CONV and IEE-FLASH (Fig. 4b). Then, the parameters were modified, as follows: a) average dose rate ; 0.2 Gy/s, dose in the bunch ; 20 mGy and dose rate in the bunch ; 6.4x10 Gy/s and b) average dose rate = 10 Gy/s, dose in the bunch = 336 mGy and dose rate in the bunch ; 10^11^Gy/s (Table S7 a&b, supplementary data). Interestingly with these parameters, a differential effect was produced, as the length deficit D10 for a) was 45% and for b) 30% (Fig. 4c) in the range of what was observed with the FLASH-IEE and a similar DMF of ∼1.5. **To our knowledge, these results are the first to show that a differential effect on ZFE can be induced by variations in bunch intensity with VHEE operating at ultra-high dose rate**, results that further define the FLASH sparing parameters at CLEAR.

To support the idea that intense electron delivery impacts biological outcome, we returned to the eRT6. The eRT6 is a versatile device but unlike the CLEAR, bunches cannot be adjusted and are similar when conventional and FLASH dose rates are used. However, variations of the average dose rate, dose rate in the pulse, dose in the pulse, pulse width, and frequency were implemented, while irradiating ZFE at 4hpf at a constant delivered dose of 10 Gy. A dose-rate escalation study was done with values ranging from 0.1 to 5.6x10^6^ Gy/s which is the maximum dose rate possible with eRT6 (Table S8, supplementary data). These experiments identified an average dose rate of 100 Gy/s, using 10 pulses of 1 Gy at 100Hz with a pulse width of 1.8 μs i.e., dose rate in the pulse of 0.5x10^6^Gy/s as threshold parameters for protecting ZFE morphogenesis (Fig. 4d). Modification of the pulse width from 1.8 μs to 4 μs when a dose of 10 Gy was delivered in one, two and five pulses (Table S9, supplementary data), did not modify the sparing outcome (Fig. 4e). However, consistently with the results obtained at CLEAR, modification of the pulse interval from 10 milliseconds C100Hz) up to 10 _min_ (Table S10, supplementary data), keeping the pulse as intense as possible, did not modify the sparing outcome (Fig. 4f). These results show that neither overall time nor the pulse number (if less than 6 pulses) are driving the biological response whereas high pulse intensity at ultra-high dose rate does trigger the FLASH sparing effect with the eRT6.

When the results obtained at eRT6 and CLEAR were combined and for a dose of 10 Gy, an additional notion emerged: for a dose rate in the range of 10^6^ Gy/s, a dose of at least 1 Gy in the pulse was needed to preserve ZFE morphogenesis but at higher dose rate, in the range of 10^11^ Gy/s, a dose in the bunch of 336 mGy was sufficient. This last result suggests that higher is the dose rate, lower the dose in the pulse/bunch can be.

### 3.6. Radiolytic H_2_O_2_ production in water and morphogenesis in ZFE are not correlated

Lastly, the FLASH parameters defined at CLEAR were used to measure H_2_O_2_ production (Table S11, supplementary data) but similar ratio were found as shown (Supp Fig. 5, supplementary data). Finally, no correlation between H_2_O_2_ production and *in vivo* results obtained in ZFE was found (Fig. 5).

**Fig. 5.**
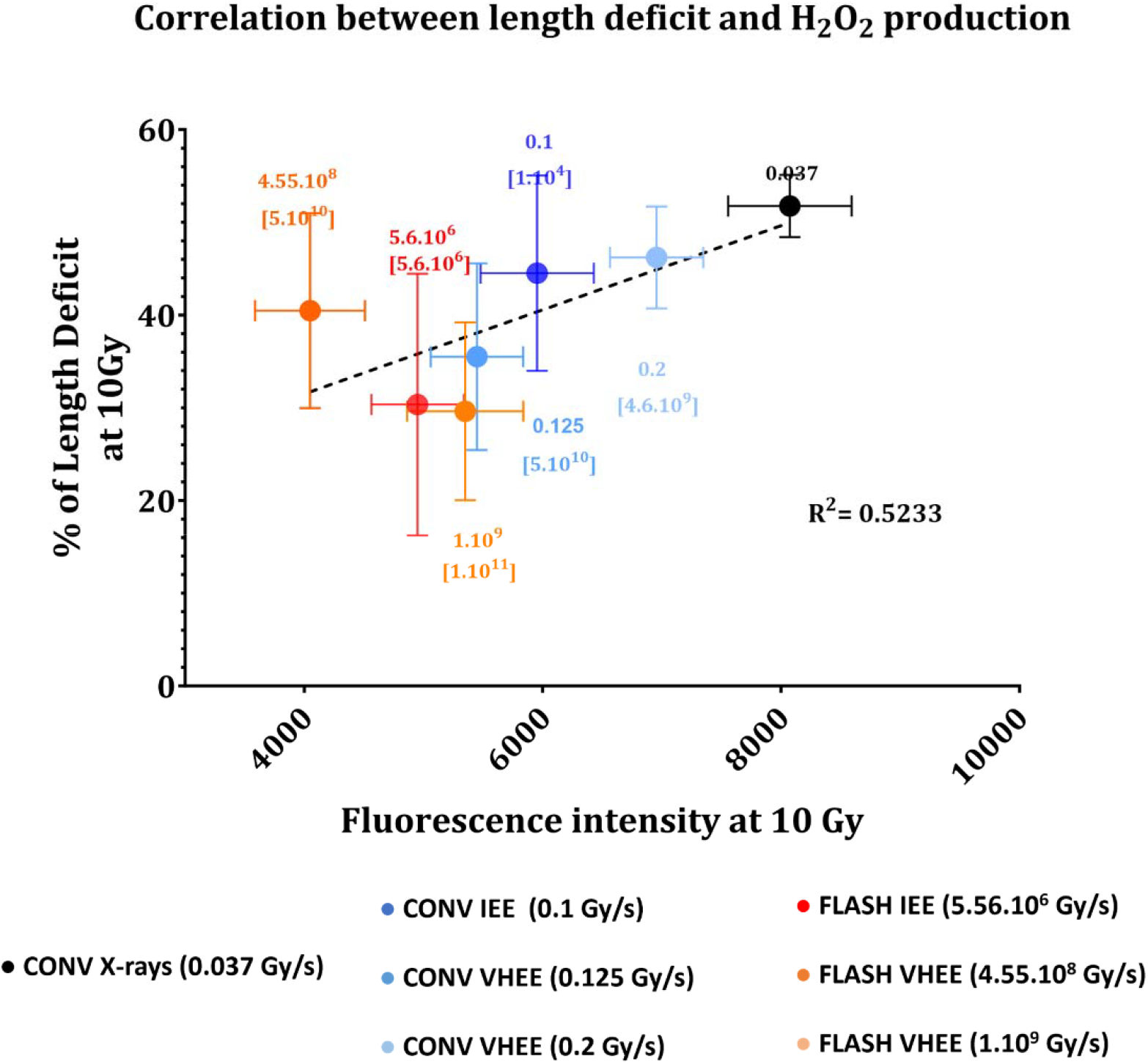
Correlation of ZFE Length deficiency and production. The percentage of length deficit after 10 Gy irradiation of ZFE represented as a function of physioxic fluorescence intensity ( production) after 10 Gy irradiation delivered with CONV and FLASH VHEE, IEE and CONV X-rays. The dashed line represents the linear fitting. The R squared = 0.5233, suggesting a poor correlation between the two models. The filled circles indicate the different modalities of beams with their average dose rates and instantaneous dose rates in brackets.

## 4. Discussion

This study is the first comprehensive exploration investigating the impact of a large range of dose rates and various temporal parameters from early physico-chemical events to a complex biological system. Our results show that reductionist radiochemical approaches using water and plasmid are poor surrogate of the *in vivo* response that define the FLASH effect. However, they show that a different biological response can be expressed days after irradiation when electrons are delivered at the pico/nano/microsecond time scale versus minutes. Importantly, we identified that the combination of high intensity bunches and ultra-high average dose rate are the critical parameters to trigger the FLASH sparing effect at CLEAR, supported by the results obtained at eRT6 with high intensity pulses. These conclusions have significant implications for designing future FLASH devices and/or guiding research into the mechanisms of FLASH.

In the present study and aligning with historical data from the water radiolysis literature, we observed that primary radiolytic yields of hydrogen peroxide remain unaffected by modulation of dose rate. These findings agree with previous measurements made using pulsed electrons [34] or conventional g-ray sources [29–31]. Additional insights can be taken from our study, namely a lower primary yield of H_2_O_2_ was found for VHEE beams as compared with IEE beams. This phenomenon may be attributed to a hypothetical overlap of spurs at high dose rates, such as the ones provided by CLEAR. In addition, the reduced secondary production of H_2_O_2_ following FLASH irradiation, also suggests a possible alteration in the subsequent chemical reactions after the homogeneous phase. These results align with our previous findings and other recent reports [35–37], yet they contradict the findings of Anderson et al. [34] and Sehested et al. [38]. These researchers reported an increased hydrogen peroxide production with pulsed electrons (> 5x10^6^ Gy/s) compared to conventional γ-rays. These discrepancies may stem from oxygen conditions. Anderson et al. and Sehested et al. used an oxygen concentration equal to 1x10^-3^ M, which is four times higher than the atmospheric oxygen pressure (2.5x10^-4^M) used in our experiments. Another point of discrepancy is brought by simulation studies and related to the increased H_2_O_2_ production with increasing dose rate. This is inconsistent with the experimental results and suggest that other type of simulations based on molecular dynamics should also be considered [39]. Whereas in a previous work, we proposed that the reduced radiolytic production of H_2_O_2_ achieved upon FLASH irradiation, could be used as a proxy to probe FLASH beam capability and even proposed the existence of a threshold, where H_2_O_2_ yield ≥ 2.33 molecules/100 could be correlated with the preservation of ZFE morphogenesis [36]. The current results invalidate this idea, as the chemical measurements in this work did not correlate with biological outcomes *in vivo* using the ZFE model, even under biochemically/biologically relevant conditions (Fig. 5).

In order to investigate more complex macromolecules, pBR322 plasmids were then irradiated. Results show that DNA damage in cell-free systems was dose-rate insensitive and not modified when beam characteristics, such as the mean dose rate, the dose rate in the bunch/pulse, beam structure, particle energy were modulated. The dose-rate independence of DNA damage in plasmids is maintained in biochemical conditions that mimic certain aspects of the tumor microenvironment (hypoxia, high Fe^2+^ levels). This highlights the insufficiency of the model to fully capture the complexity of biological *in vivo* models. Our results are consistent with earlier results by Milligan et al. [40,41] performed with dose rates ranging between 0.1 Gy/s and 1 Gy/s that included scavengers, as well as with published results produced at CLEAR [42] with dose rates above 10 Gy/s. However, they contradict a recent report claiming reduced DNA damage in plasmids using UHDR electrons (46.6 Gy/s) [43]. In this later study, experimental conditions were probably improperly controlled as the plasmid was not properly purified which likely confounded their conclusions. In addition, data was fitted based on a very low number of experimental data points that further confounded a rigorous interpretation. However, in a recent study Wanstall et al. [44] irradiated pBR322 plasmids with VHEE CLEAR using two sets of beam parameters comparable to those applied in the current study involving ZFE. Interestingly, variation of the bunch dose rate and intensity induced measurable difference in single strand breaks at doses above 90 Gy. Those results are in line with the findings of the current work. The relevance of DNA damage after FLASH *vs* CONV has been explored *in vivo* in mice and lead to contradictory conclusions. Fouillade et al. [45] reported less gH2AX foci after FLASH electrons (2x10^2^ to 4x10^7^Gy/s) in the normal lung, Levy et al. [46] showed similar levels of γH2AX foci in ID8 tumors after FLASH electrons (216 Gy/s) and CONV electrons (0.079 Gy/s). They also showed a slight decrease in repair foci after FLASH in the normal gut at early time-points, that similar residual foci by 24h post-irradiation [46]. In an *ex-vivo* study of whole-blood peripheral blood lymphocytes (WB-PBL), using a comet assay, Cooper et al. [47] also reported reduced DNA damage after electron FLASH (2000 Gy/s) at doses higher than 20 Gy and under reduced oxygen conditions (0.25 − 0.5 % O_2_). Barghougth et al. [48] used a functional and quantitative DNA damage repair (DDR) assay *in vitro* and showed that electron FLASH (1x10^2^ to 5x10^6^Gy/s) did not alter chromosome translocations and junction structures in HEK293T cells more than CONV electron dose rates (0.08 Gy/s to 0.13 Gy/s) did. Collectively, the sum of experimental data obtained so far suggest that DNA damage is unlikely to explain the FLASH effect and results obtained with plasmids suggest that DNA damage is clearly dose rate independent at lower doses (below 10 Gy) and might be dose-rate dependent at high doses (above 20 Gy).

Our most important finding was that living organisms are sensitive to variation of electron beam intensity. At CLEAR, high intensity bunches of electrons (336 mGy) at ultra-high average dose rate (average: 10^9^ and instantaneous: 10^11^ Gy/s) spares ZFE morphogenesis while lower intensity bunches (20 mGy) at ultra-high average dose rate (average: 0,2 and Instantaneous: 6.4.10^9^ Gy/s) are more toxic to ZFE. Bunches of intermediate intensity (150 mGy) but delivered at a different average dose rate (0.125 vs 4.5510^8^ Gy/s) lead to the same length deficit. A similar trend is observed with the eRT6 when high intensity pulses of 1 Gy or more are delivered at high average dose rate (5.6.10^6^ Gy/s) corresponding to the microsecond scale. These data indicate that intensity is a key parameter. In addition, the higher the dose rate, the lower the dose needed to trigger the FLASH sparing effect (i.e., 0.3 Gy at 10^11^ Gy/s and 1 Gy at 10^6^ Gy/s). This is another critical observation, as it helps establish the therapeutic basis of operating FLASH in the clinical setting, where small doses per fraction remain standard of care. Interestingly, this observation is supported by results from the literature where morphogenesis sparing required doses above 30 Gy with dose rates ranging from 300 Gy/s to 7500 Gy/s in ZFE irradiated 24 hpf [49]. Furthermore, Ruan et al. [50] showed as well that the highest number of intestinal crypt cells in young mice was spared with a dose of 11.2 Gy delivered in a single FLASH pulse of 3.4 μs of electron irradiation. When the number of pulses and the time interval between two pulses (i.e., the frequency) was increased, crypt cells number was still maintained when compared to CONV [50]. The idea that high intensity pulses trigger the best dose modifying factor has also been recently confirmed by Liu et al. in mouse. Fr a dose of 12 Gy, when they increased the dose per pulse (DPP) from 1 to 6.1 Gy with 1.5-1.7 MGy/s in the pulse and an average dose rate of 130-188 Gy/s, enhanced preservation of the number of regenerating crypts was shown. In addition, for the same when 2 pulses of 6 Gy were spaced by 8.33 millisec to 34 sec at 1.7 MGy/s, crypt protection was sustained. Interestingly, these conditions remained efficient against melanoma [51]. Our observations also show that the sparing of living organisms (at least ZFE) is not likely to be enhanced further at higher intensity and dose rate and suggest that the benefits of normal tissue protection reach a plateau above 10^6^ Gy/s.

## 5. Conclusion

In conclusion, we provide the first evidence that live organisms are sensitive to variations of electron beam intensity, where compression of beam delivery time can impact biological response. While these observations are both transformational and challenging to traditional radiobiology, they do provide an evidence-based framework for the design of future FLASH clinical accelerators. Present results suggest that pencil/spot scanning techniques used to cover target volumes are feasible and FLASH compatible with VHEE and proton beams. Given the reduced doses required for FLASH sparing at high intensities and dose rates, VHEE beams seem optimally poised for clinical developments of FLASH radiotherapy.

## Credit authorship contribution statement

**Vozenin Marie-Catherine** Conceptualization – Investigation – Supervision – Writing original draft – Securing fundings. **Houda Kacem** Investigation – Methodology – Writing original draft. **Louis Kunz** Investigation – Methodology – Writing original draft. **Pierre Korysko** Investigation – Methodology – Dosimetry. **Jonathan Ollivier** Investigation – Methodology. **Pelagia Tsoutsou** Writing original draft. **Adrien Martinotti** Investigation. **Vilde Rieker** Dosimetry. **Joseph Bateman** Dosimetry. **Wilfrid Farabolini** Investigation – Dosimetry. **Gérard Baldacchino** Simulation. **Billy W. Loo Jr** Writing original draft. **Charles L. Limoli** Conceptualization – Supervision – Writing original draft. **Manjit Dosanjh** Conceptualization – Supervision – Dosimetry. **Roberto Corsini** Conceptualization – Supervision – Dosimetry. **All authors** Write, Review & Edit.

## Acknowledgments

The authors thank Dr C Bailat, Prs F. Amati, J. Bourhis, F. Bochud for their support. We also thank the PTZ-UNIL.

## Supplementary Materials

### Supplementary Text

#### Dosimetry

The doses delivered during sample irradiations using VHEE beams at the CLEAR Facility were measured using radiochromic films. Gafchromic EBT3 and MD-V3 films (Ashland Inc., Bridgewater, NJ, USA) were used for doses up to 10 Gy, and above 10 Gy, respectively. On sample holder, two radiochromic films were positioned in front and behind of the PCR tube, separated by 12 mm. The radiochromic films were scanned with an Epson Perfection V800 Photo scanner (Epson, Long Beach, US) at least 24 h after irradiation. Dose-to-water calibrations for both film types were performed at CONV dose rates on the eRT6 at CHUV. For each point on the calibration curve the dose was measured using a PTW Advanced Markus Ionisation Chamber (PTW, Freiburg, Germany). The radiochromic film analysis was performed following the single channel method outlined in work by Micke *et al*. (2011) [52] and is similar to that used for previous UHDR studies using radiochromic film dosimetry at the eRT6 (Jaccard *et al* 2017) [53]. A lead pellet which occupies the same volume as the sample is placed inside an additional PCR tube holder with a radiochromic film located behind the PCR tube. This creates a resulting shadow on the dose distribution of this radiochromic film, from which an area of interest (AOI) can be delineated and applied to the radiochromic films used for dosimetry of the irradiated samples. The mean and standard deviation of the dose across this AOI is calculated for both films on each holder, and the dose to the sample is determined as the average of this mean dose across both films. The beam size is calculated by applying a Gaussian fit to the dose distribution and obtaining the standard deviation of the Gaussian fit in both the *x* and *y* directions. Work by Rieker *et al*. (2023) [54] describes some of the initial radiochromic film and passive dosimetry studies with VHEE beams at the CLEAR Facility.

#### Water radiolysis experiments

##### Aqueous solutions

Milli-Q water was used with a conductivity of 18.2 µS/cm. For scavenging experiments, aqueous solutions were prepared of different concentrations of NaNO_2_ from 10 μM to 100 mM and one constant concentration of [NaNO_2_] = 25 mM. Water samples were equilibrated in glass bottles at room temperature in hypoxia hood (Biospherix) to achieve hypoxic oxygen conditions (1% O_2_) for 48h. The day of the experiment, water was transferred to required tubes and irradiated to 10 − 50 Gy with the different beams. Hydrogen peroxide production at 21% O_2_ was performed in Milli-Q water without any scavenger and irradiated to similar doses as for the scavenging experiments.

##### Measurement of the irradiated samples

Water samples were probed immediately after irradiation with Amplex Red assay kit purchased from Thermo Fisher. Amplex Red was added at a final concentration of 50 µM and incubated for 30 min protected from light. Freshly H_2_O_2_ solutions from 0.3125 μM to 10 μM were prepared and used to establish the calibration curve. Fluorescence quantification was performed using Promega Glo-Max plate reader (Excitation: 520 nm Emission: 580 − 640 nm). G-value of hydrogen peroxide was calculated from the slope of plots of hydrogen peroxide concentrations as a function of the irradiated dose.

**Fig. S1.**
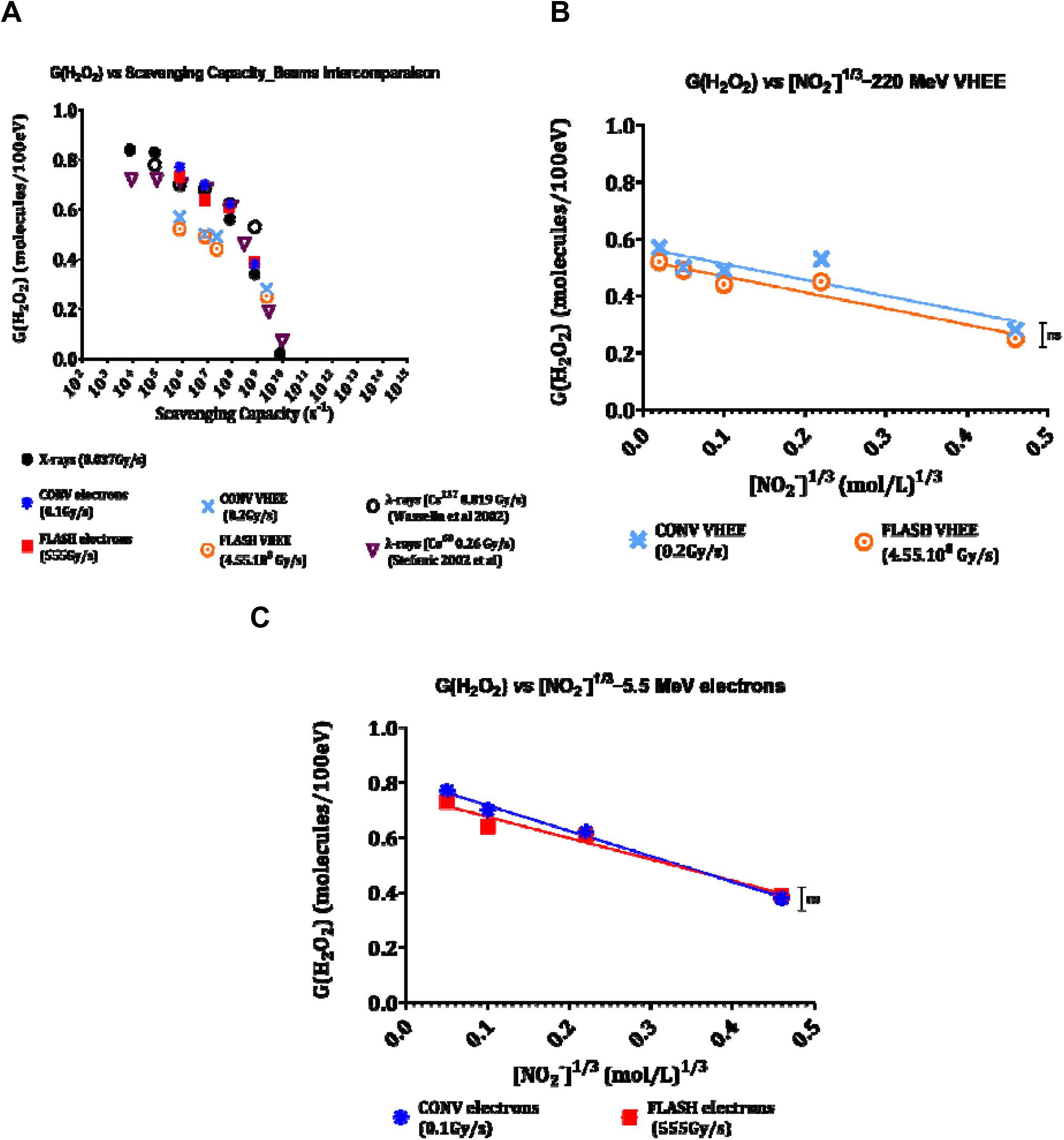
Primary hydrogen peroxide yields. **A** G-value of hydrogen peroxide yields as a function of the scavenging capacity of a scavenger ( ions for this work) obtained after irradiation with CONV and FLASH VHEE, IEE and CONV X-rays and compared to reported irradiations with CONV ^137^Cs g-rays; [S] = ], ( ) and CONV ^60^Co g-rays; [S] = [ , ( ). **B**) production from solutions of various concentrations and constant concentrations of after exposure to VHEE at CONV and FLASH. **C**) production from solutions of various concentrations and constant concentrations of after exposure IEE at CONV and FLASH.

**Fig. S2.**
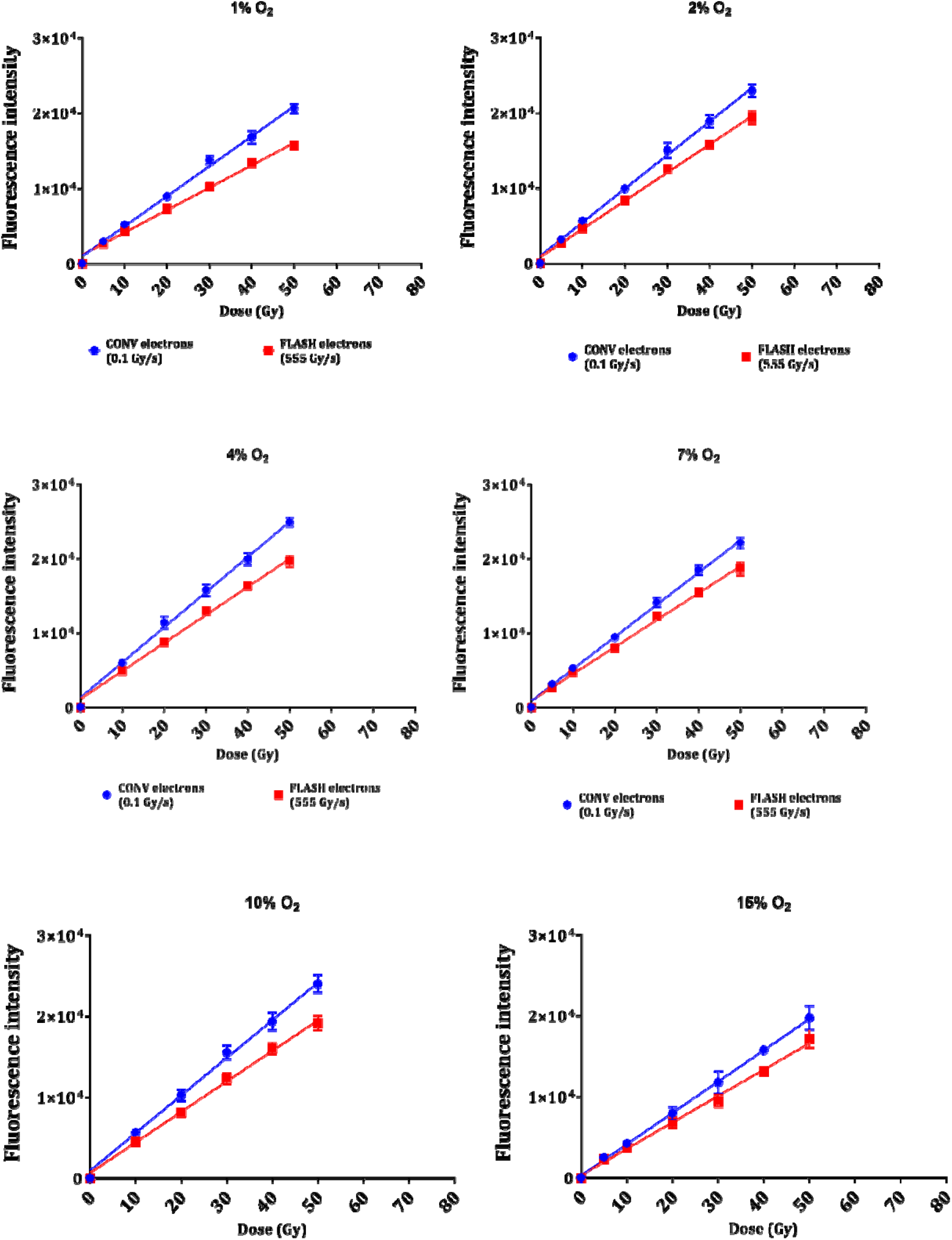
Impact of oxygen on hydrogen peroxide production. Production of hydrogen peroxide as a function of the dose after irradiation with CONV ( ) and FLASH ( ) IIE at different oxygen levels. Experiments are results of duplicate irradiation.

**Fig. S3.**
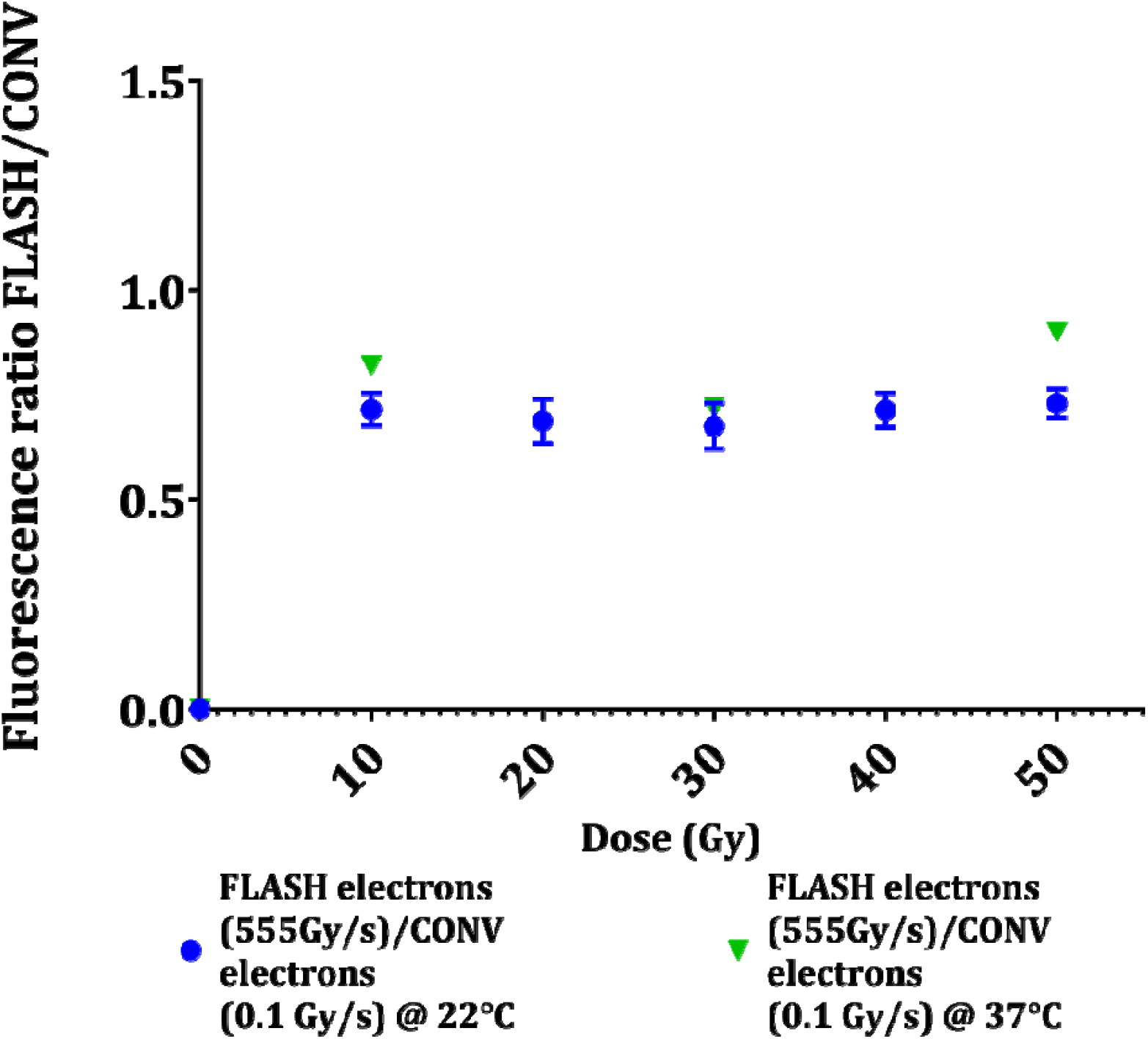
Impact of temperature on hydrogen peroxide production. Fluorescence ratio FLASH/CONV at room temperature ( ) and physiological temperature ( ) after exposure to CONV ( ) and FLASH ( ) IEE. Experiments are results of triplicate irradiations.

**Fig. S4.**
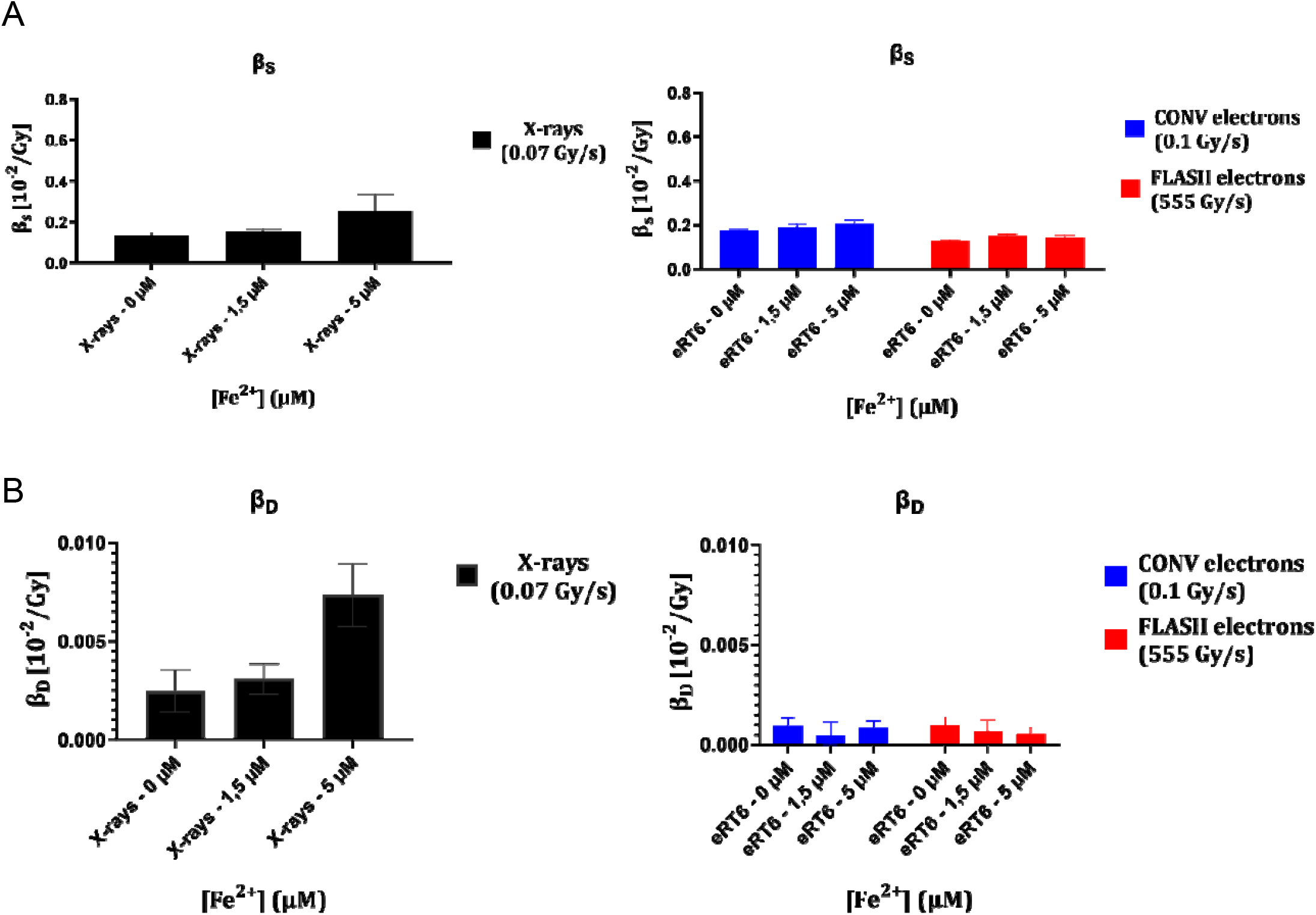
DNA damage yields after CONV X-rays and IEE (CONV and FLASH) irradiation. **A** (SSB yields) and **B** (DSB yields) of DNA damage in plasmids obtained in hypoxic conditions ( ) and diluted with different ( ) concentrations ( ) obtained after exposure to CONV X-rays ( ) and CONV ( ) and FLASH ( ) IEE.

**Fig. S5.**
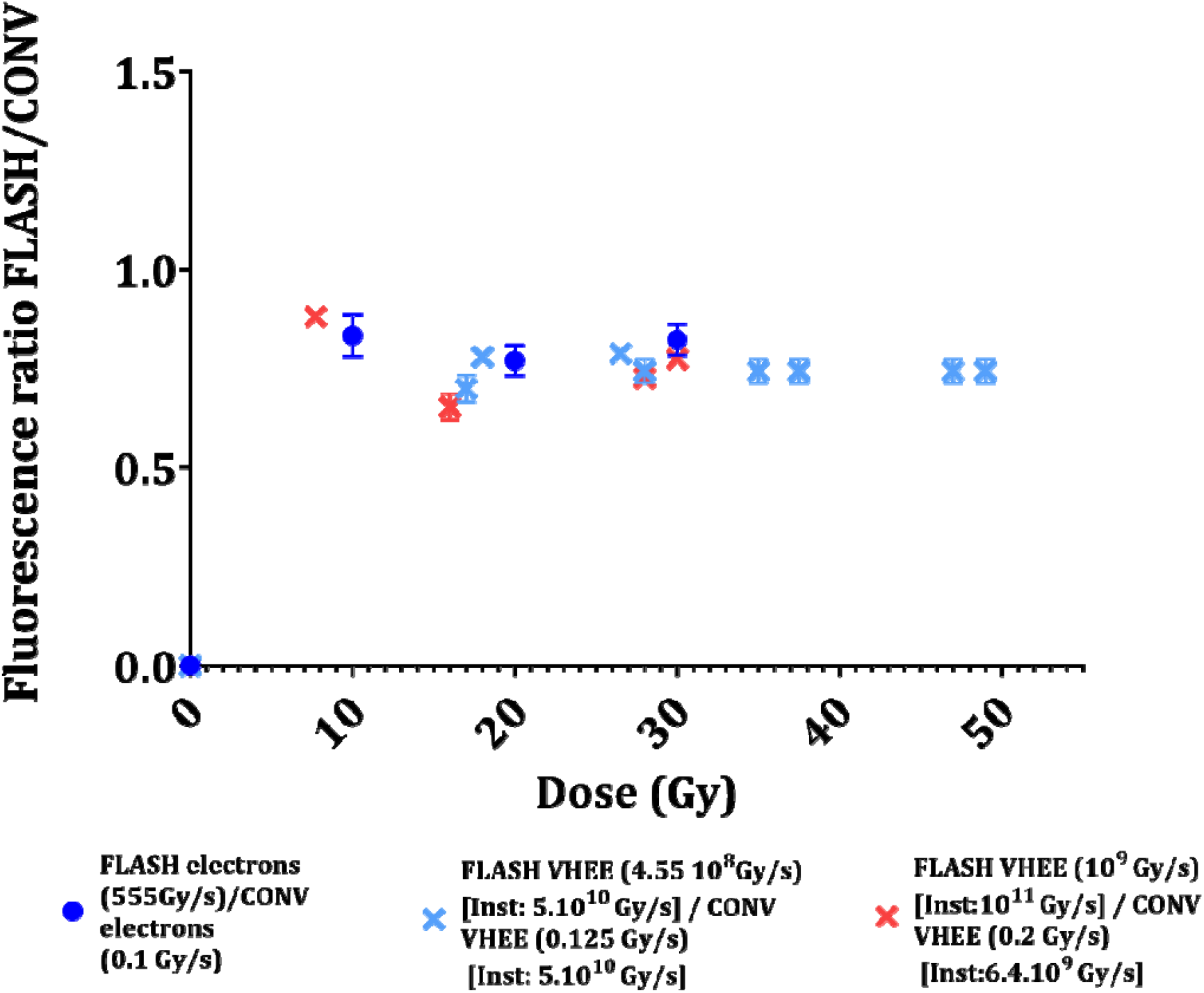
Impact of modifying CLEAR beam parameters on hydrogen peroxide production. Fluorescence ratio FLASH/CONV production *vs* dose in pure water after irradiation with CONV and FLASH IEE, VHEE with similar and different instantaneous dose rate in the bunch respectively.

